# Luminal breast epithelial cells from wildtype and *BRCA* mutation carriers harbor copy number alterations commonly associated with breast cancer

**DOI:** 10.1101/2024.05.01.591587

**Authors:** Marc J. Williams, Michael UJ Oliphant, Vinci Au, Cathy Liu, Caroline Baril, Ciara O’Flanagan, Daniel Lai, Sean Beatty, Michael Van Vliet, Jacky CH Yiu, Lauren O’Connor, Walter L Goh, Alicia Pollaci, Adam C. Weiner, Diljot Grewal, Andrew McPherson, McKenna Moore, Vikas Prabhakar, Shailesh Agarwal, Judy E. Garber, Deborah Dillon, Sohrab P. Shah, Joan Brugge, Samuel Aparicio

## Abstract

Cancer-associated mutations have been documented in normal tissues, but the prevalence and nature of somatic copy number alterations and their role in tumor initiation and evolution is not well understood. Here, using single cell DNA sequencing, we describe the landscape of CNAs in >42,000 breast epithelial cells from women with normal or high risk of developing breast cancer. Accumulation of individual cells with one or two of a specific subset of CNAs (e.g. 1q gain and 16q, 22q, 7q, and 10q loss) is detectable in almost all breast tissues and, in those from *BRCA1* or *BRCA2* mutations carriers, occurs prior to loss of heterozygosity (LOH) of the wildtype alleles. These CNAs, which are among the most common associated with ductal carcinoma in situ (DCIS) and malignant breast tumors, are enriched almost exclusively in luminal cells not basal myoepithelial cells. Allele-specific analysis of the enriched CNAs reveals that each allele was independently altered, demonstrating convergent evolution of these CNAs in an individual breast. Tissues from *BRCA1* or *BRCA2* mutation carriers contain a small percentage of cells with extreme aneuploidy, featuring loss of *TP53*, LOH of *BRCA1* or *BRCA2*, and multiple breast cancer-associated CNAs in addition to one or more of the common CNAs in 1q, 10q or 16q. Notably, cells with intermediate levels of CNAs are not detected, arguing against a stepwise gradual accumulation of CNAs. Overall, our findings demonstrate that chromosomal alterations in normal breast epithelium partially mirror those of established cancer genomes and are chromosome- and cell lineage-specific.

## Introduction

Somatic mutations are known to accumulate in normal tissues over time and, although the vast majority are inconsequential, contribute to cancer^1–3^. Most studies have measured and emphasized the role of single nucleotide variants (SNVs) in normal tissues. Yet gene dosage mutations due to somatic copy number alterations occur in the majority of tumor types^4,5^ and are highly prevalent in breast cancers^6–8^, contributing important driver events such as *ERBB2* amplification and *PTEN* loss. They also represent the dominant source of transcriptional variation in genomically unstable human cancers^4,6,9–11^, including breast cancer. Studies of pre-invasive DCIS have noted that extensive CNAs and structural variants (SV), resulting from duplication or loss of whole chromosome or chromosome segments, are already present with a landscape largely indistinguishable from invasive cancers^12,13^. Early pre-cancer atypical ductal hyperplasias are also noted to have extensive CNA mutations^14,15^. These findings indicate that CNAs arise early in the evolution of breast cancer; however, a full understanding of the prevalence, evolutionary timing and distribution of the earliest CNAs arising in morphologically normal breast epithelium is lacking.

The vast majority of SNV mutations are private to single cells or form small clonal expansions that would be obscured by bulk short read sequencing of tissues. We posit this is also the case for CNAs. Recent studies of SNVs in normal tissues have successfully used a combination of ultra-deep error corrected sequencing^16^ or experimental cloning amplification of single cells subsequently characterized with bulk short read next generation sequencing^17,18^ to bypass these barriers. However, the prevalence of CNAs in most normal cells may be an order of magnitude or more lower than SNVs and thus comprehensive characterization of CNAs is inaccessible to these approaches. A few studies have attempted to discover somatic CNAs in normal tissues^19–23^ by reanalyzing bulk sequencing data but have been limited to blood or to detecting CNAs present in >20% of the cellular population, which do not allow the underlying generative process of CNAs in individual cells to be defined. We have overcome these limitations by developing methods for scaled single cell whole genome sequencing (scWGS) (DLP+)^24,25^ which allow for discovery of CNAs unique to single cells in thousands of individual genomes. By sampling without restriction directly from tissues, the progeny of single mitotic mutational events leading to cell-specific alterations can be ascertained.

Here we investigate the prevalence and landscape of copy number alterations in normal breast epithelial tissues to identify the earliest genetic alterations using DLP+ scWGS. We reveal the prevalence, chromosomal distribution, and lineage specificity of CNA mutations in breast tissues from high risk *BRCA1*/*BRCA2* germline mutation carriers and contrast with BRCA-wildtype epithelium.

## Results

### Low aneuploidy prevalence in normal mammary epithelia is cell type dependent

To assess the distribution and prevalence of CNAs in single breast epithelial cells of individuals with germline breast cancer predisposition alleles, we obtained breast tissues from women carrying germline pathogenic mutations in *BRCA1* (n=8) and *BRCA2* (n=6) undergoing risk-reducing surgery, as well as from those with the *BRCA1/2* wild-type (WT) genotype (n=6) from reductive mammoplasties. Some women had a history of breast cancer or other cancers and had received prior chemotherapy ( **Fig. 1a**). See **Supplementary Table 1** for all clinical details. For patients with a history of breast cancer, tissue was acquired from the contralateral breast. Macroscopically normal tissue was allocated for research purposes. Microscopic examination of representative FFPE blocks of clinical and/or research tissue revealed no atypical hyperplasia or in situ carcinoma in 15/20 subjects. Representative tissue samples from 5 donors revealed small foci (<1-2mm) of in situ carcinoma or atypical hyperplasia: B2-16 (DCIS), WT-7 (ADH), WT-6752 (ALH), B2-21 (ALH), B2-23 (LCIS) (Supplementary Table 1). Tissue samples were then dissociated into single cells, sorted into luminal and basal cell populations based on previously established surface markers (^26–28^ methods) and the single cell genomes sequenced to an average genome-wide coverage of 0.029X using the DLP+ protocol^29^ (range 0.001-0.361, **Supplementary Table 2**). After removing low quality genomes and discarding samples with fewer than 300 cells, 42,756 single cell genomes from 20 donors were analyzed (**Fig. 1a**). Example genome wide copy number profiles from a diploid genome and aneuploid genome are shown in **Figure 1b-c**.

**Figure 1.**
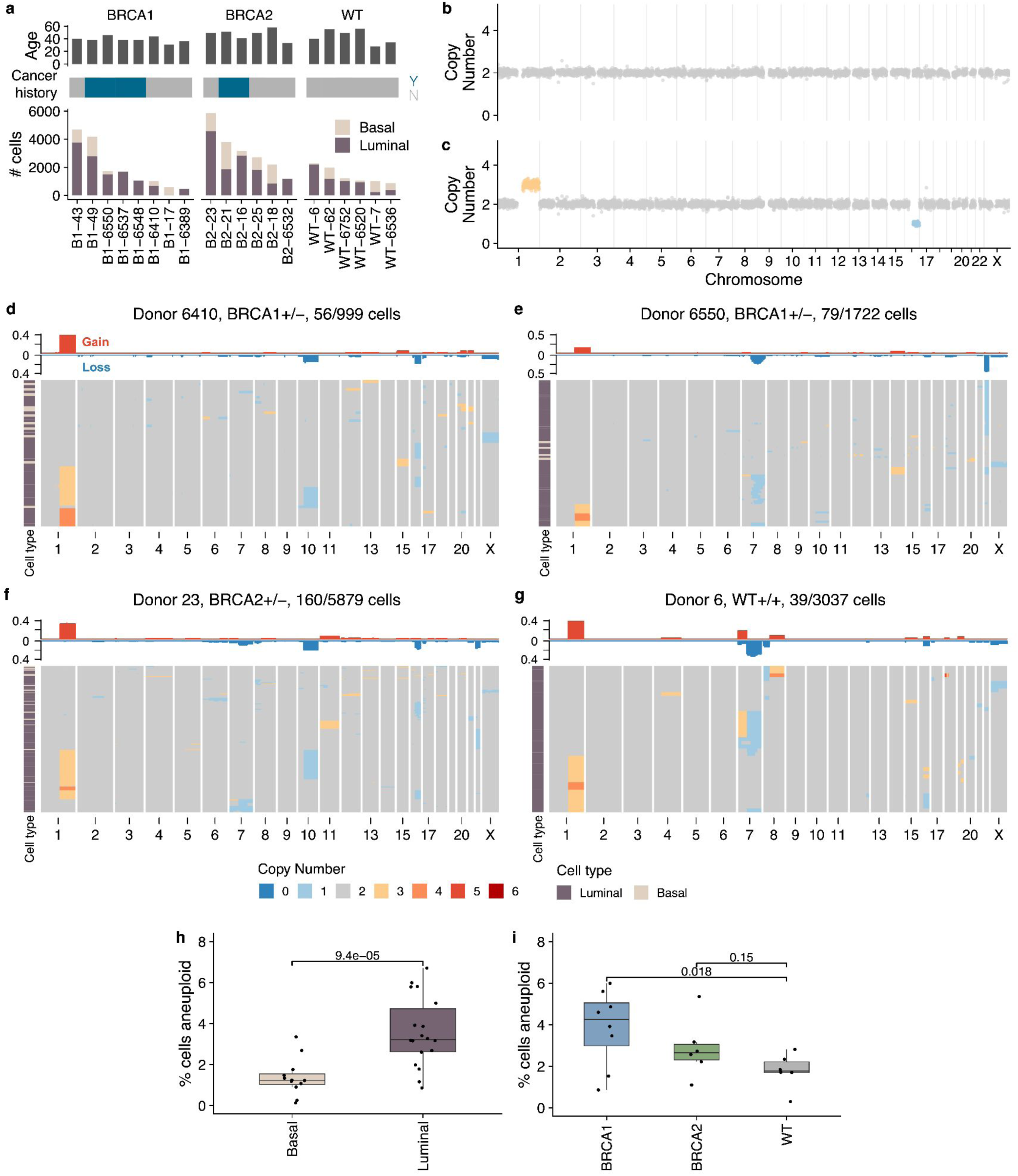
**a)** Number of high quality cells per sample per cell type along with cancer history and patient ages **b)** Example diploid cell **c)** Example aneuploid cell with chr1q gain and chr16q loss **d)** Heatmap of aneuploid cells from donor B1-6410, title shows donor name, genotype and number of aneuploid cells out of total number of cells **e)** Heatmap of aneuploid cells from donor B1-6550 **f)** Heatmap of aneuploid cells from donor B2-23 **g)** Heatmap of aneuploid cells from donor WT-6 **h)** %of cells aneuploid between cell types **i)** % of cells aneuploid between genotypes

Aneuploid cells, defined as cells with at least one chromosome arm level gain or loss, were rare but observed in every sample. Overall, 2.69% of cells (range: 0.1-5.9%) contained between one and four aneuploid chromosome arms (simple aneuploidy). Notably, specific alterations such as gains of 1q and losses on 16q, 10q, 22q and 7q were recurrent across donors for four samples: two BRCA1 ^+/-^ (B1-6410 and B1-6550), one BRCA2^+/-^ (B2-23) and one WT (WT-6) (**Figure 1d-g**). Similar patterns were observed in all other donors, see **Supplementary Figure 1a-c**. These results indicate that cells carrying a specific subset of CNAs accumulate in ostensibly normal breast epithelial cells.

Aneuploid cells were more prevalent in luminal cells compared to basal cells (3.6% vs. 1.4%, p=9.4x10^-5^, **Fig. 1h**), and in BRCA carrier donors compared to WT: 3.8% in BRCA1 and 2.9% in BRCA2 compared with 1.8% in WT donors (p=0.02 and p=0.15 respectively, **Fig. 1i**). We did not find any significant associations with other clinical covariates including age, parity, menopause status, cancer history or chemotherapy history (**Supplementary Figure 2**). In a multi-variate regression that included age, genotype and cell type, luminal cells were associated with an increase in aneuploidy (p=5.69x10 ^-5^) and the WT genotype with a decrease in aneuploidy (p=0.024); no other groups showed a statistically significant association (**Supplementary Figure 2f**).

### Recurrent aneuploidies in luminal cells are similar to breast cancers

Next, we explored the distribution of CNAs across the genome and between cell types. Luminal and basal cells had distinct distributions of CNAs. CNAs observed recurrently across patients were restricted to luminal cells (**Fig. 2a** & **Supplementary Figure 3**). These included gain of 1q, the most common observed alteration (1.06% in luminal vs 0.03% in basal, p=0.00009), loss of 16q (0.6% vs 0.04%, p=0.00044), loss of 22q (0.5% vs 0.03%, p=0.0049), loss of 7q (0.33% vs 0.01%, p=0.0025) and loss of 10q (0.27% vs 0.07%, p=0.032 **Fig. 2** & **Supplementary Figure 3**). Loss of chromosome X was also common but occurred at similar rates in both luminal and basal cell types (0.16% vs 0.12%, p=0.63, **Fig. 2a,b** & **Supplementary Figure 3**). Since X chromosome loss has been shown to increase with age and preferentially involve the inactive copy^21^, it is likely a selectively neutral event that would explain the approximately equal rate of loss in the two cell types. We did not identify any alterations that were statistically significantly more prevalent in basal cells compared to luminal cells.

**Figure 2.**
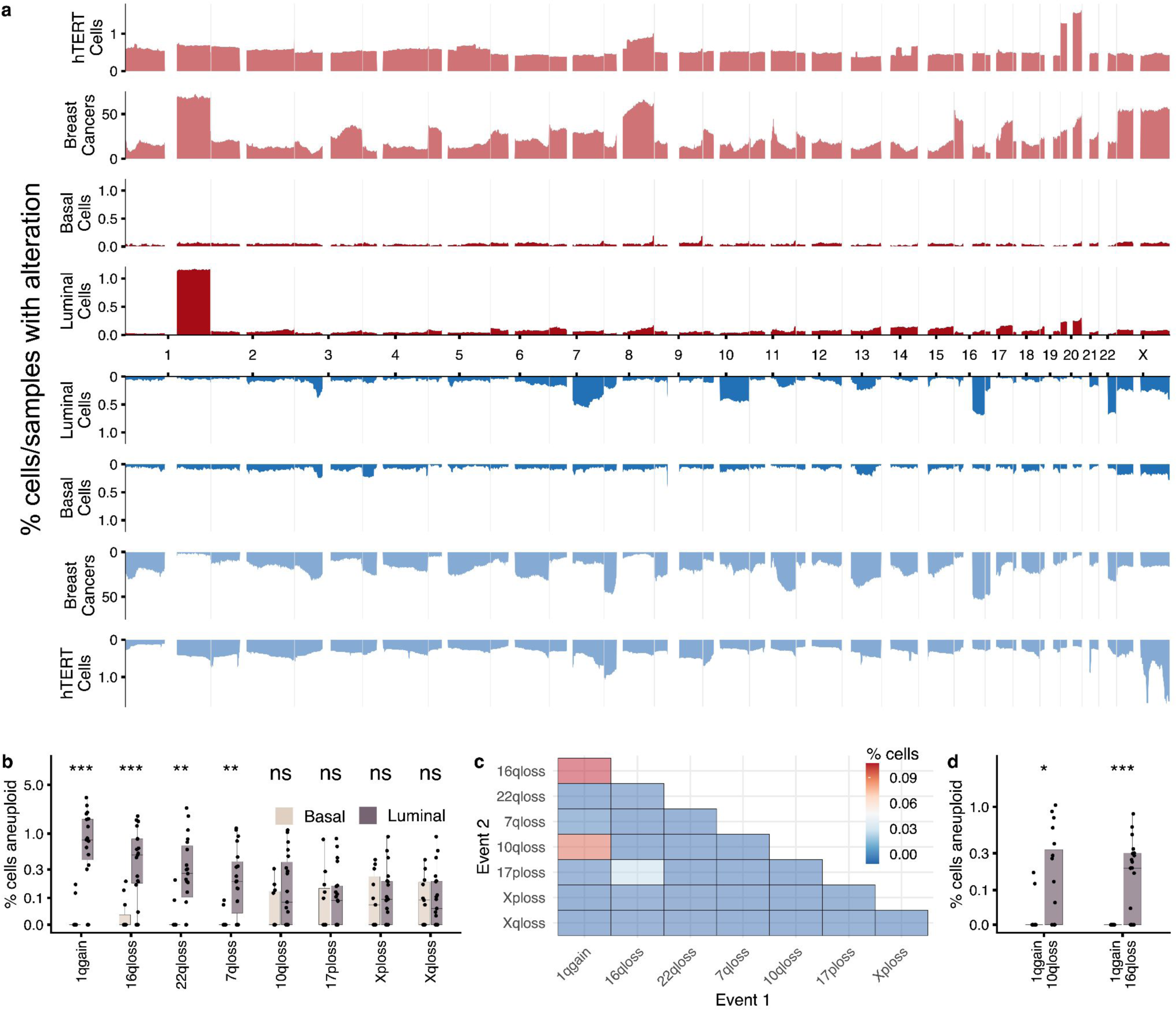
**a)** Frequency of gains/losses across the cohort, y-axis is fraction of cells or samples that have gains/losses. 3 cohorts shown. hTERT cells: 14,000 cells from an immortalized mammary epithelial cell line, Breast Cancers: 555 whole genome sequence cancers from Nik-Zainal et al. Luminal and basal cells from this study **b)** % cells aneuploid per patient split by luminal and basal cells for the 8 most common chromosome alterations **c)** co-occurence heatmap showing percentage of cells that have 2 chromosomal aneuploidies concurrently **d)** % of cells that have 1q-gain/16q-loss and 1q-gain/10q-loss per cell type

To assess how these patterns compare to those from invasive breast cancers, we compared the normal tissue CNA chromosomal distribution to 560 whole genome sequenced breast cancers from Nik-Zainal *et al*^30^. A number of events that were common in the luminal cell population were also common in advanced cancers including the gains of 1q and losses of 16q and 22q (**Fig. 2a**). Loss of 7q, which is common in our normal epithelium dataset, is comparatively rare in breast cancers (**Fig. 2a**). Conversely, there are some events such as gains of 8q and 16p and loss of 11q that are very common in breast cancers but are rare in normal breast epithelium, suggesting that these alterations are typically acquired later during tumor evolution. Computing the cosine similarity between normal tissue CNA distributions and all cancer types present in the TCGA, we found that breast cancers were the most similar cancer type for both gains and losses, (**Supplementary Figure 4**). We note the similarity to some other cancer types, which reflects the fact that some of the common alterations (e.g. 1q gain) are also prevalent in other cancer types.

To explore whether the enrichment of certain chromosomes could be explained by underlying mutational bias, we also compared the distribution of CNAs to that derived from 14,000 single cell genomes from a wild-type immortalized breast tissue cell line (hTERT cells). In contrast to the scWGS from normal breast epithelium, the distribution of CNAs in this cell line was relatively uniform across the genome ( **Fig. 2a**). This suggests that chromosome arms have a relatively uniform susceptibility to CNAs and that the higher prevalence of CNAs within certain chromosomes in normal breast epithelium is a tissue- and cell type-specific process, potentially linked to lineage differentiation and/or epithelial cell orientation within a tissue context^31^.

Amongst cells that had more than one aneuploid chromosome arm, the most frequent events were 1q-gain/16q-loss (present in 12 donors) and 1q-gain/10q-loss (present in seven donors, **Fig. 2c**). Both combinations were enriched in luminal cells with average frequencies of 0.23% (1q-gain/16q-loss) and 0.19% (1q-gain/10q-loss,**Fig. 2d**). Interestingly, 10q-loss was only ever observed in conjunction with 1q-gain while 16q-loss was frequently observed in isolation. These data are consistent with a recent report^32^ that showed that clones carrying 1q-gain/16q-loss events are precursors that emerge decades before cancer diagnosis.

### Allele-specific alterations reveal multiple independent CNAs

To address whether the recurrent aneuploidies that we observed arose from single clonal expansions or constituted multiple independent events, we phased chromosome gains and losses to parental alleles (here defined arbitrarily as allele A or B) using SIGNALS^33^, a HMM based inference approach determining allele-specific copy number alterations. Observing gains and losses of both alleles would indicate that these events had been acquired independently more than once and give a lower bound on the number of events.

Applying SIGNALS to 10 samples that contained a large number of aneuploid cells, we found evidence that CNAs were independently acquired at least twice. For example, B2-23 had aneuploid cells with all the frequent CNAs: 1q-gain, 7q-loss, 10q-loss, 16q-loss and 22q-loss and also several cells with both 1q-gain/10q-loss and 1q-gain/16q-loss (**Fig. 3a**). Allele-specific copy number analysis revealed gains and losses on each allele, indicating each event must have been acquired independently at least twice (**Fig. 3b**). In the case of cells with 1q-gain/10q-loss, we could infer three separate configurations: 1q(A-gain)-10q(B-loss), 1q(B-gain)-10q(B-loss) and 1q(B-gain)-10q(A-loss) (**Fig. 3b**). Similarly, for cells with 1q-gain/16q-loss, most had lost the B-allele on 16q but we identified one cell that had lost the A-allele.

**Figure 3.**
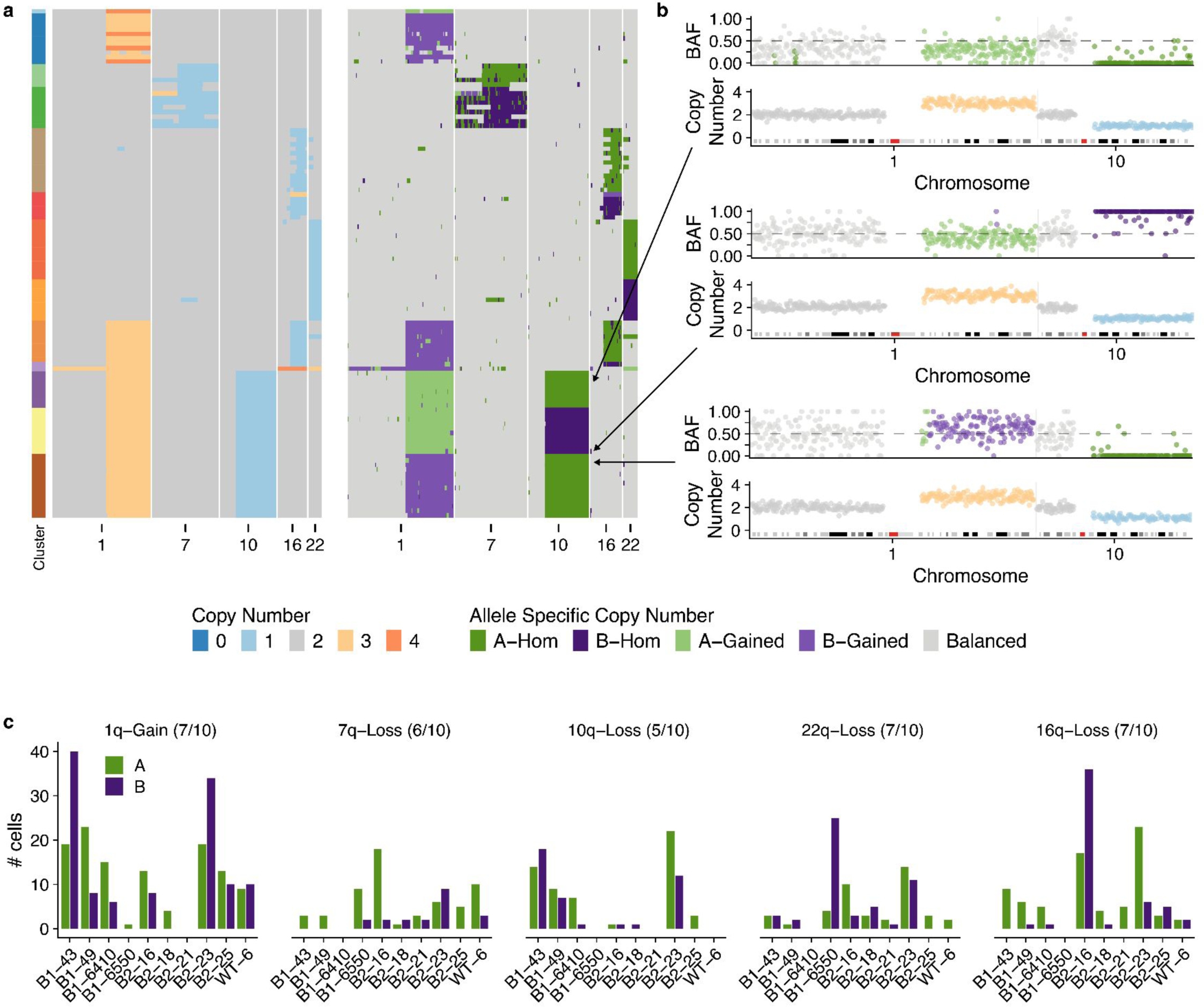
**a)** Total copy number heatmap and allele specific copy number heatmap for B2-23 for chromosomes 1,7,19,16 & 1. 22. Cells grouped into unique alterations based on allele specific copy number. Total number of cells = 111 **b)** Three cells from the heatmap with chr1q gain and chr10q loss. For each cell the B-allele frequency BAF and copy number is shown for chromosomes 1 and 10. These 3 cells have distinct combinations of chr1-gain and 10 loss. **c)** Number of cells with either allele A or B gained/lost across the 6 most common alterations in 10 patients. Title above each plot shows the event and the number of samples that have events on both alleles

Applying the same analysis to an additional nine samples, we found that there was evidence that the common alterations were acquired independently multiple times in the majority of cases. For example, cells with gain of 1q of both alleles were present in 7/10 samples, and losses of both alleles on 7q and 16q were observed in 6/10 and 7/10 samples, respectively. Taken together, these findings indicate that the aneuploid populations we observe are not part of a single clonal expansion but rather are consistent with multiple independent alterations, all of which are able to survive and proliferate. Furthermore, this also suggests alterations on either allele have similar phenotypic effects.

### Extreme aneuploid cells are rare but present across individuals

Some models of cancer evolution posit that highly aneuploid genomes of invasive breast cancers could emerge from single catastrophic mitosis with multiple chromosomal defects as opposed to progressive accumulation of events over multiple mitoses^34^. To shed light on this, we searched for cells with extreme aneuploidy. The majority of aneuploid cells have at most one or two CNAs, however, there exists a small population of cells with many CNAs (**Fig. 4a**). We classified extreme aneuploid cells as those exceeding 9 aneuploid chromosome arms, placing them in the upper 5% of the CNA burden distribution ( **Fig. 4a**). Extreme aneuploid cells were rare but present across individuals with an average prevalence of 0.1% (range 0-0.43%) (**Fig. 4b** & **Supplementary Figure 5** for heatmaps). We then calculated how similar these single cell genomes were to the average breast cancer profile and identified 23 cells that were similar (ρ≥0.25), labeling these “cancer-like” genomes (**Fig. 4c**).

**Figure 4.**
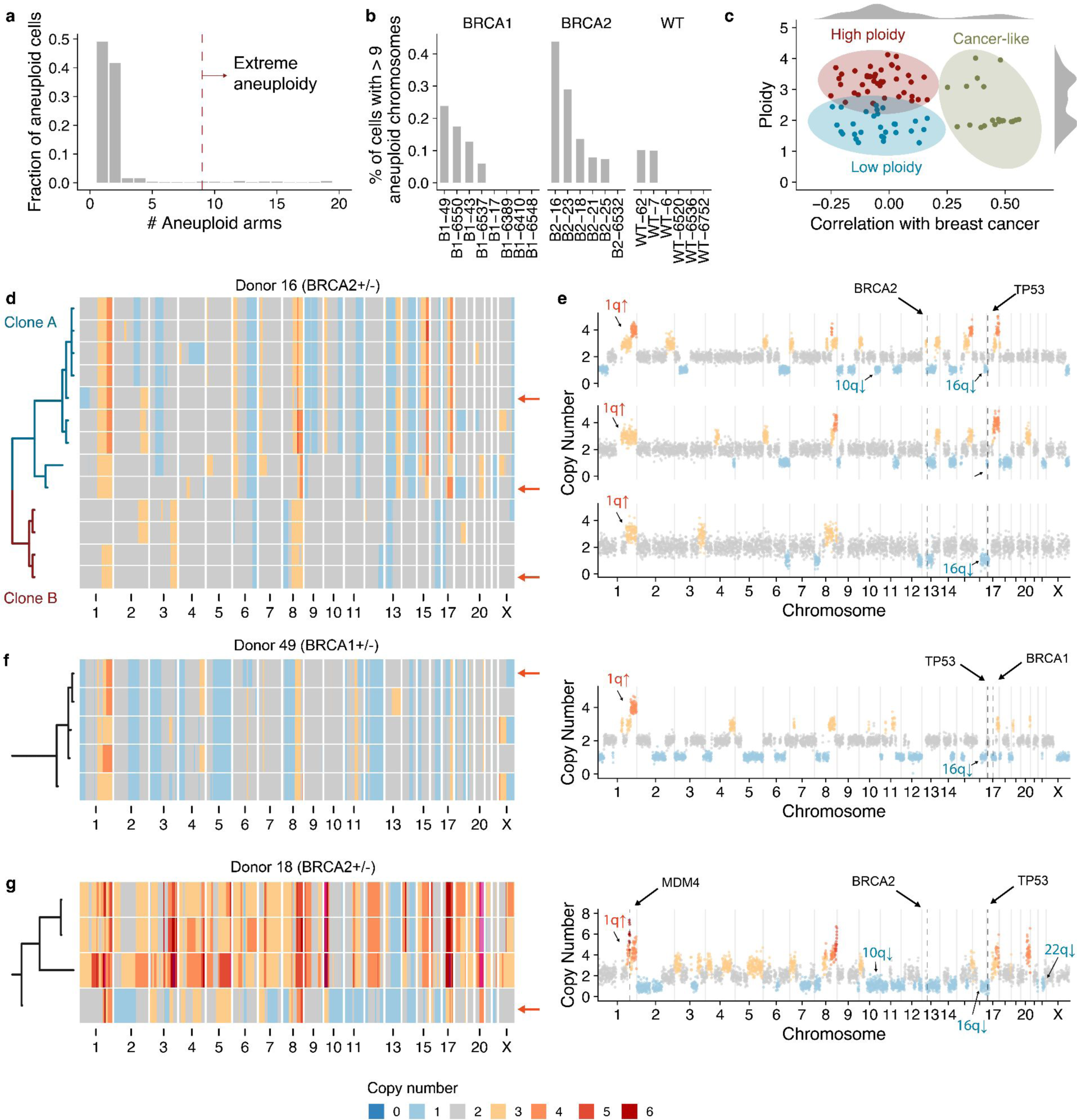
**a)** Fraction of the aneuploid cells that have X aneuploid arms. Dashed red line shows the cutoff (=9) used to classify cells as extreme aneuploidy **b)** % of cells in each sample with > 9 aneuploid chromosomes. **c)** Scatter plot of ploidy vs correlation with cancers from Nik-Zainal *et al.* highlighting three distinct groups: high ploidy, low ploidy and cancer-like **d)** Heatmap of extreme aneuploid cancer-like cells in patient B2-16 ordered by a phylogenetic tree. **e)** 3 cells from patient 16 with arrows showing their placement in the heatmap. **f)** Example cell and heatmap of extreme aneuploid cancer-like cells in patient B1-49 **g)** Example cell and heatmap of extreme aneuploid cancer-like cells in patient B2-18

The 23 “cancer-like” cells were derived from three high-risk donor samples. All “cancer-like” cells had lost one copy of either *BRCA1* or *BRCA2*, although we cannot be certain that the wild-type copy was lost due to the inability to confirm mutational status in individual cells due to the limited sequencing coverage per cell. All cells had also lost one allele on 17p, the location of *TP53*, suggesting that these cells had also lost P53 function. B2-16 has 13 cancer-like cells that through phylogenetic analysis could be subdivided into two independent clones, clone A and clone B (**Fig. 4d,e**). Although both these clones share similar features such as gains on 1q and 8q and losses on 6q, 16q, 13p (including *BRCA2*) and 17p (including *TP53*), the copy number changepoints for these events are distinct in each clone, strongly suggesting they are evolutionary independent clonal lineages. This is further supported by allele-specific analysis showing different alleles lost in chromosomes 6 and 16 in the two clones (**Supplementary Figure 6a**).

B1-49 had five “cancer-like” cells that were all evolutionary related (**Fig. 4f**). All cells had gains of 1q and 8q, and losses on 16q and 17q (including *BRCA1*). Allele-specific analysis also revealed that 17p was copy neutral LOH (**Supplementary Figure 6b**). B2-18 had four “cancer-like” cells that again, were all evolutionary related (**Fig. 4g**). These cells had gains on 1q, 8q and 17q and losses on 10q, 13q (including *BRCA2*), 17p (including *TP53*), 16q and 22q among others. Interestingly 3/4 cells had undergone a whole genome doubling, while one cell – that likely resembles the ancestral state of the three other cells – remained in a diploid state. Pathological review of these breast tissues revealed a small DCIS lesion associated with one of the FFPE blocks of B2-16.

We note that in samples with these cancer-like genomes, we did not observe cells with intermediate aneuploid states that might be expected from a stepwise gradual accumulation of CNAs. This could reflect the possibility that intermediate states are unfavorable to cellular proliferation or cleared by immune cells or, alternatively, that all the changes are acquired within a short period of time, or plausibly a single mitotic event.

Amongst the cells that were not correlated with advanced breast cancers (ρ<0.25) (**Fig. 4c**), a significant proportion were characterized by a large number of whole chromosome losses relative to cell ploidy (see **Supplementary Figure 5** & **Supplementary Figure 7**). These cells are consistent with cytokinesis failure or multipolar divisions and are likely non-viable as we rarely observed two cells with near identical genomes. Furthermore, in some cases, such cells had large regions that were homozygously deleted (**Supplementary Figure 7**). However, there was a notable example of a clonally expanded genome doubled population (n=14 cells) in donor B2-23 (**Supplementary Figure 5**).

## Discussion

This study of scaled single cell genome analysis of breast epithelium reveals several striking features of somatic copy number alterations in pathologically normal tissues. First, we show that aneuploidy is uncommon, comprising 2.69% overall of epithelial cells. Second, we observe a marked difference in epithelial lineages: luminal cells, the putative precursor compartment for breast malignancies, exhibit 3.6% aneuploid cells, whereas only 1.4% of basal myoepithelial cells carried aneuploidies. Third, we observed that CNAs occur with structured tissue architecture across the genome: the most abundant CNAs were largely limited to the luminal population and included gains on 1q and losses on 10q, 16q, 22q and 7q. Loss of chromosome X was similar in luminal and basal lineages, which may be explained by the loss of the inactive copy being selectively neutral. Fourth, this specific pattern of CNAs may be tissue context specific, as we did not observe it in cultured mammary epithelial cells. Thus, our data suggests that CNAs form a significant component of the somatic mutational spectrum of epithelial cells in normal breast tissues, and this is both chromosome- and cell lineage-specific, even within mammary epithelial sub-lineages.

When compiling individual CNA events across many single genomes into an aggregate, the normal cell CNA landscape we observe bears a striking resemblance to bulk sequencing data of invasive breast cancers. One of the most commonly observed alterations from our dataset was co-occurring 1q gain and 16q loss in luminal epithelial cells. Interestingly, these co-occurring CNAs are often found to be the only alteration present in low grade DCIS and luminal A tumors^7,35,36^. Our data not only support that concurrent 1q gain and 16q loss is an early event, but that it is almost exclusively associated with luminal epithelial cells and can occur through multiple independent allelic events. Concurrent 1q-gain/16q-loss is most often generated through an unbalanced translocation event that results in the fusion of chromosome 1q and 16p arms, termed der(1;16)^37,38^. Interestingly, a recent phylogenetic analysis identified der(1;16) as a founder alteration that could be traced back to early pubertal breast epithelial cells. These clones expanded over time and acquired additional mutations that eventually led to cancer development^32^ (**Supplementary Figure 8**). While 1q/16q CNAs were found to be the only CNAs for some low grade tumors, these alterations are also associated with high aneuploid tumors^38^. Due to limitations in the resolution of our sequencing data, we were unable to confirm whether 1q-gain/16q-loss clones in our dataset were a result of der(1;16). Nevertheless, our results strongly support the importance of premalignant alterations in 1q and 16q and raise the question whether targeting of early progenitors harboring 1q-gain/16q-loss may be an effective therapeutic strategy for preventing or monitoring breast cancer development.

While 1q gain as the most commonly detected event, additional alterations were repeatedly identified including co-occurring 1q gain and 10q loss, 7q loss, and 22q loss. All of these CNAs, with the exception of 7q loss, are enriched in breast tumors. Although these alterations occurred at lower prevalence, some have been implicated as predictive of subtype and prognosis^6,7,36,39^. For example, 10q loss is of particular interest because *PTEN* is located on this chromosome arm and deletions of *PTEN* are commonly associated with basal breast tumors (TCGA). *PTEN* loss has also been computationally predicted to occur prior to *BRCA1* LOH in human breast tumors^40^.

We speculate the CNA mutational events that accumulate later in the progression from normal epithelium to cancer may be dependent on these earlier alterations. For example, it is known that MYC overexpression sensitizes cells to apoptosis and survival of high MYC cells requires anti-apoptotic alterations like p53 loss of function or gain of BCL2 anti-apoptotic proteins ^41–43^. The *MDM4* suppressor of p53 is on 1q and 1q gain in tumor cells has been shown to increase the expression of MDM4, suppress p53 signaling, and is associated with *TP53* mutations that are mutually-exclusive with 1q aneuploidy in human cancers^44^. The anti-apoptotic protein MCL1 is also located on 1q. Thus, it is possible that CNAs are required to tolerate significant alterations as cells undergo transformation. Notably, some common breast cancer associated CNAs such as 8q are not prevalent in mammary epithelium, suggesting these are selected later in cancer evolution.

In addition to the cells with one or two CNAs, we also detected a small number of cells in *BRCA1* and *BRCA2* mutation carriers with extensive CNAs, which were similar to those that occur in BRCA-mutant cancers^45,46^. These cells may derive from microscopic pre-malignant lesions present in the donor tissue. Most of these cells also carried CNAs in 1q and 10q or 16q, raising the possibility that the presumed loss of the WT *BRCA* allele occurred in cells with the pre-existing CNAs. It is of interest that we did not observe an intermediate set of alterations progressing from minimal to extreme aneuploidy. The paucity of intermediate clones in our analysis supports a punctuated model of clonal evolution, which proposes tumor development as abrupt transitions rather than a gradual accumulation of alterations over time^47,48^. Therefore, we hypothesize (**Supplementary Fig 8**) that cells with minimal aneuploidy may serve as founder cells that undergo rapid bursts of alterations triggered by catastrophic events like LOH of *BRCA1* or *BRCA2*, TP53 loss of function, chromothripsis or whole-genome duplication. Alternatively, intermediate states may be more susceptible to immune surveillance leading to rapid elimination or require additional alterations to overcome LOH and undergo transformation. These intriguing hypotheses require further investigation, with longitudinal studies potentially shedding light on the dynamics of clonal evolution of cells with CNAs, as well as providing additional insights into the relationship between cancer-associated genetic alterations and immune activity during early stages of tumorigenesis.

The patterns we observe could be due to a mutational bias (e.g. preferential mis-segregation of certain chromosomes^49^, contribution of chromosome specific fragile sites) or differing relative fitness of cells carrying CNAs. Although the sampling method used here captures the single cell background, largely bypassing purifying selection and not reliant on clonal amplification for detection of CNAs, measuring actual contributions of potential hypermutability and/or fitness to the landscape would require the timing and population fitness of individual CNAs to be measured. This is not currently tractable from human tissues at single cell resolution. Nevertheless, taken together, our data suggest that the mechanisms of somatic copy number alterations and/or selection operate continuously in non-malignant epithelium, emphasizing the need to better understand the mechanistic relationships between lineage specific mutational and selection forces in tumor formation.

## AUTHOR CONTRIBUTIONS

JSB and SA conceived this study. MJW, MUJO, JSB and SA wrote the manuscript with input from other authors. MJW analyzed all scDNAseq data. MUJO organized tissue sample processing, dissociated and processed tissues, and carried out FACS sorting. LO dissociated and processed tissues, WG processed and FACS-sorted samples. JEG, DAD, AP, MM, and orchestrated tissue procurement. Shailesh Agarwal and ACP acquired patient consent, VP performed tissue collection and initial processing after surgery, DAD performed pathological reviews. SPS supervised computational analysis. DL, CL, SB, DG, AM, AW, JCHL developed and ran computational pipelines. VA generated the scDNAseq data with support from CO’F, MVV and CB.

## Acknowledgements

We gratefully acknowledge the teams who facilitated tissue collection for these studies, including the BWH breast surgery team led by Dr. Tari King; the BWH plastic surgery team; and the BWH Faulkner pathologists and technical staff led by Dr. Tony Guidi. We thank Ron Schackmann, Abdu Alsaadi, Gianmarco Rinaldi, Kung-Chi Chang, Kate Moore and Klarisa Norton for support in processing tissues. We also thank the DFCI Flow Cytometry Core led by John Daley and Suzan Lazo. We deeply appreciate invaluable editorial feedback provided by Drs. M. Angelica Martinez-Gakidis (JSB lab). This work was supported in part by a Gray Foundation Team Sciences Award (JSB, SA, DAD, JEG,), a Goldberg Family Research Fund gift (JSB), the Breast Cancer Research Foundation (JSB), an Anbinder Cancer Research Fund gift (JSB), and an NCI grant NCI R35 CA242428(JSB). MJW is supported by a National Cancer Institute Pathway to Independence award (K99CA256508). MUJO is supported by the R35 Diversity Supplement (R35CA242428-04) and the Black in Cancer/Emerald Foundation Inc Postdoctoral Career Transition Fellowship. S.P.S. holds the Nicholls Biondi Chair in Computational Oncology and is a Susan G. Komen Scholar (GC233085). S.A. holds the Nan and Lorraine Robertson Chair in Breast Cancer and is a Canada Research Chair in Molecular Oncology (950–230610). Additional funding was provided by a Terry Fox Research Institute grant (1082), CIHR grants (FDN-148429, 495630), Breast Cancer Research Foundation awards (BCRF-21-180, BCRF22-180, BCRF23-180) and the Canada Foundation for Innovation (40044) to S.A.

## Tables

**Supplementary Table 1**

Clinical details of the 20 donor patients including BRCA1/2 mutations, age, cancer history, chemotherapy history, details on pathological review, parity and menopause status

**Supplementary Table 2**

Cell level statistics including cell_id, sample, cell_type, cell coverage, number of aneuploid arms and extreme aneuploidy classification.

## Methods

### Tissue procurement

All donor samples analyzed in the study are listed in Table S1. Specimens were obtained from Brigham & Women’s Hospital or Faulkner Hospital on the day of surgery. This study was reviewed by the Harvard Medical School Institutional Review Board (IRB) and deemed not human subjects research. Donors gave their informed consent to have their anonymized tissues used for scientific research purposes. The scDNAseq dataset contains 20 samples that include 6 elective reduction mammoplasties and 14 prophylactic mastectomies (7 *BRCA1* mutation carriers, 6 *BRCA2* mutation carriers and 1 *BRCA1/BRCA2* mutation carrier). The age range of the cohort is 28-58 years old.

### Tissue processing and FACS

Breast tissue samples were dissociated as previously described^50^. Briefly, each tissue was minced and transferred to a 50 ml conical tube containing a solution of Advanced DMEM/F12 (Thermo 12634010), 1× Glutamax (Gibco 35050), 10 mM HEPES (Gibco 15630), 50 U/ml Penicillin-Streptomycin (Gibco 15070) and 1 mg/ml collagenase (Sigma C9407). Digestion was performed by constant shaking at ∼150-200 rpm at 37C for 2-4 hours. Tissue was then pelleted by centrifugation and further dissociated into single cells by treatment with TrypLE (Gibco 12605010) for 5-15 min. After neutralization and pelleting by centrifugation, sequential pipetting with 25, 10 and 5 ml pipette tips was performed to further dissociate the tissue. The dissociated tissue was then filtered through a 100um and 40um filter to isolate single cells and counted manually under the microscope to assess yield and viability. Single cells were fixed with 1.6% paraformaldehyde for 10 min and cryopreserved until ready for FACS.

For FACS isolation of mammary epithelial cell types, single cells isolated from tissue were labeled for 30 min at room temperature with Alexa Fluor 647-conjugated anti-EpCAM (1:50, Biolegend 324212), PE-conjugated anti-CD49f (1:100, Biolegend 313612), FITC-conjugated anti-CD31 (1:100, Biolegend 303103) and Alexa Fluor 488 anti-CD45 (1:100, Biolegend 304017). The lineage-negative population was defined as CD31^-^ CD45^-^. After staining, FACS was performed to isolate CD31/CD45^-^ EpCAM^+^ CD49f^+/-^ (Luminal) and CD31/CD45^-^ EpCAM^low^ CD49f^+^ (Basal/myoepithelial) cells for scDNAseq analysis.

### Single cell DNA sequencing

We used the DLP+ protocol to generate low pass whole genome sequencing data^24^. Frozen single-cells were thawed, washed and pelleted in DMEM (Corning 10-013-CV) and resuspended in PBS (Corning 21-040-CV) with 0.04% BSA (Cedarlane 001-000-162). Single-cell suspensions were labeled with CellTrace CFSE dye (ThermoFisher C34554) and LIVE/DEAD Fixable Red stain (ThermoFisher L23102) by incubation at 37°C for 20 min. Cells were resuspended in PBS with 0.04% BSA and aspirated into a contactless piezoelectric dispenser (Scienion CellenOne) for single cell dispensing into open nanowell arrays (TakaraBio SmartChip) preprinted with unique custom dual indexed sequencing primers. Nanowell chips were subsequently scanned on a Nikon TI-E inverted fluorescent microscope (10X magnification). Singly-occupied wells and cell state were determined using our custom image analysis software, SmartChipApp (Java) (Laks et al. 2019). Cell-spotted nanowell chips are covered with SmartChip Intermediate Film (Takara 430-000104-10) and stored at -20°C until library construction.

Lysis buffer comprised of 6.73 nL DirectPCR Lysis Reagent (Viagen 302-C), 2.69 nL protease (Qiagen 19155), 0.5 nL glycerol (100%), and 0.09 nL pluronic (10%) were dispensed into each well. Nanowell chips were sealed with Microseal A (BioRad MSA5001) using a pneumatic sealer and centrifuged before each incubation step. Cells were allowed to soak overnight in lysis buffer for 18-19 hours at 21°C (30°C lid) in a flatbed thermocycler (ThermoFisher ProFlex Dual Flat PCR System 4484078). Following overnight presoak, chips were incubated at 50°C for 1 hour to carry out thermal and enzymatic lysis. Lysis inactivation (75°C for 15 min, 10°C forever) was conducted after lysis. Tagmentation was performed with 7.5 nL Bead-Linked Transposomes (BLT, Illumina DNA Prep 20060059), 7.5 nL Tagmentation Buffer 1 (TB1, Illumina DNA Prep 20060059), and 15 nL nuclease-free water, incubated at 55°C for 15 min. Neutralization was carried out with 9.9 nL protease (Qiagen 19155) with 0.1 nL Tween20 (10%) at 50°C for 15 min, followed by heat inactivation at 70°C for 15 min. Limited-cycle PCR amplification was conducted with 44.53 nL Enhanced PCR Mix (EPM, Illumina DNA Prep 20060059) and 0.47 nL Tween20 (10%) using the following conditions: 68°C for 3 min; 98°C for 3 min; 11-cycles of 98°C for 45 sec, 62°C for 30 sec, 68°C for 2 min; 68°C for 1 min; and hold at 10°C. Single-cell whole genome libraries were eluted from nanowell chips by centrifugation through a funnel into a recovery tube. Pooled libraries were cleaned by double-sided bead purification using Sample Purification Beads (SPB, Illumina DNA Prep 20060059) and eluted into Resuspension Buffer (RSB, Illumina DNA Prep 20060059).

Single-cell whole genome libraries were quantified with Qubit dsDNA High Sensitivity Assay (ThermoFisher Q32854) and Bioanalyzer 2100 HS kit (Agilent 5067-4626). Sequencing was conducted to a depth of 0.03X coverage per cell on either: Illumina NextSeq 2000 (2x100 bp) at UBC Biomedical Research Centre (Vancouver, BC), Illumina HiSeq 2500 (2x150 bp) or Illumina NovaSeq 6000 (2x150 bp) at the BC Genome Sciences Centre (Vancouver, BC).

### Single cell DNA processing and analysis

The single cell-pipeline outlined in Laks *et al.* was used to call copy number in single cells at 0.5Mb resolution. Briefly, this pipeline aligns sequencing reads to the reference genome, counts the number of reads in 0.5Mb bins across the genome, performs GC correction using a modal regression framework and then computes integer copy number states across the genome using HMMcopy^51^. We then applied the cell quality filter and removed cells with quality < 0.75. In addition, to remove possible low quality cells not captured by the cell quality score, cells undergoing replication and cells with possible incorrect ploidy estimates we also removed cells that had the following characteristics: i) ploidy > 5 ii) >10 segments with size <5Mb.

We computed allele-specific copy number for the aneuploid cells using SIGNALS for 10 donors. As input, SIGNALS requires haplotype block counts per cell which in turn requires identifying heterozygous SNPs and phased haplotype blocks. To identify heterozygous SNPs, all cells were merged into a single pseudobulk bam file and treated as a normal whole genome sequencing sample. The “Haplotype Calling” submodule (step 8: https://github.com/shahcompbio/single_cell_pipeline/blob/master/docs/source/index.md) was then used to infer haplotype blocks and genotype them in single cells. These results were then used in SIGNALS with default parameters apart from *mincells* which was set to 4. *mincells* is the size of the smallest cluster used to phase haplotype blocks, and needed to be lower than what is typically recommended for cancer data due to the sparsity of CNAs. Downstream analysis and all plotting was done using SIGNALS^33^.

### Aneuploidy in single cells

Single cells were called as aneuploid if they had at least one chromosome arm in a copy number state that was different from the ploidy of the cell. Integer cell ploidy was assigned to be the most common copy number state across the whole genome (unless this was 1, in which case ploidy was set to 2) and chromosome arm copy number states in each cell were assigned based on the most common copy number state of the bins within a chromosome arm (using per_chrarm_cn function in SIGNALS). Aneuploid arms with copy number states greater than cell ploidy were classed as gains and less than cell ploidy as losses. Cells were classed as “Extreme Aneuploid” if they were in the top 5% of cells in terms of CNA abundance. This cutoff corresponded to 9 or more aneuploid arms.

### Additional datasets used in this study

To compare the distribution of CNAs to cancer cells we made use of whole genome sequencing data from Nik-Zainal et al^30^ and SNP array data from TCGA^10^. To facilitate comparison with scWGS DLP data, the various formats used in these studies were converted into a format that consisted of integer copy number at 0.5Mb across the genome. Gains and losses were defined relative to cell ploidy as for the single cell data.

We also used a set of >14,000 human telomerase reverse transcriptase (hTERT) immortalized wild-type mammary epithelial cells. Details of culture conditions can be found in Funnell *et al*^52^.

### Classifying extreme aneuploid cells

For each extreme aneuploid cell we computed its correlation coefficient with the average copy number profile from 262 cancer samples that had purity > 0.5 in Nik-Zainal et al. Plotting the distribution of correlation coefficients we observed a bimodal distribution, with a mode at 0, a mode at ∼0.5 and an inflection point at 0.25. We therefore classified cells that had ≥ 0.25 correlation coefficient as “cancer-like” and those with correlation < 0.25 as low ploidy or high ploidy depending on their cell ploidy, which also exhibited a bimodal distribution.

### Phylogenetic trees

We constructed phylogenetic trees for the cancer-like extreme aneuploid cells using sitka^52^ which uses copy number changepoints as phylogenetic markers. Here, a copy number change point is the locus (bin) where the inferred integer copy number state changes between bin *i* and bin *i+1*. The input to sitka is a binary matrix consisting of cells by changepoint bins. Default parameters were used. Length of branches in the trees represent the number of copy number changes.

### Statistical analysis

For between group comparisons we used t-tests. To investigate multiple factors that might influence aneuploidy while taking into account that most donors have basal and luminal cells we performed a multi-level multivariate model (**Supplementary Figure 2f**) that included cell type, age and donor genotype. We used the lmer package in R with the following formula specification: percentage_aneuploidy ∼ age + cell_type + genotype + (1|sample).

## Data availability

Raw sequencing data will be available from EGA under accession EGAS00001007716 at the time of publication.

## Code availability

Single-cell pipeline for processing DLP+ data is available at https://github.com/shahcompbio/single_cell_pipeline.

---

**Supplementary Figure 1a.**
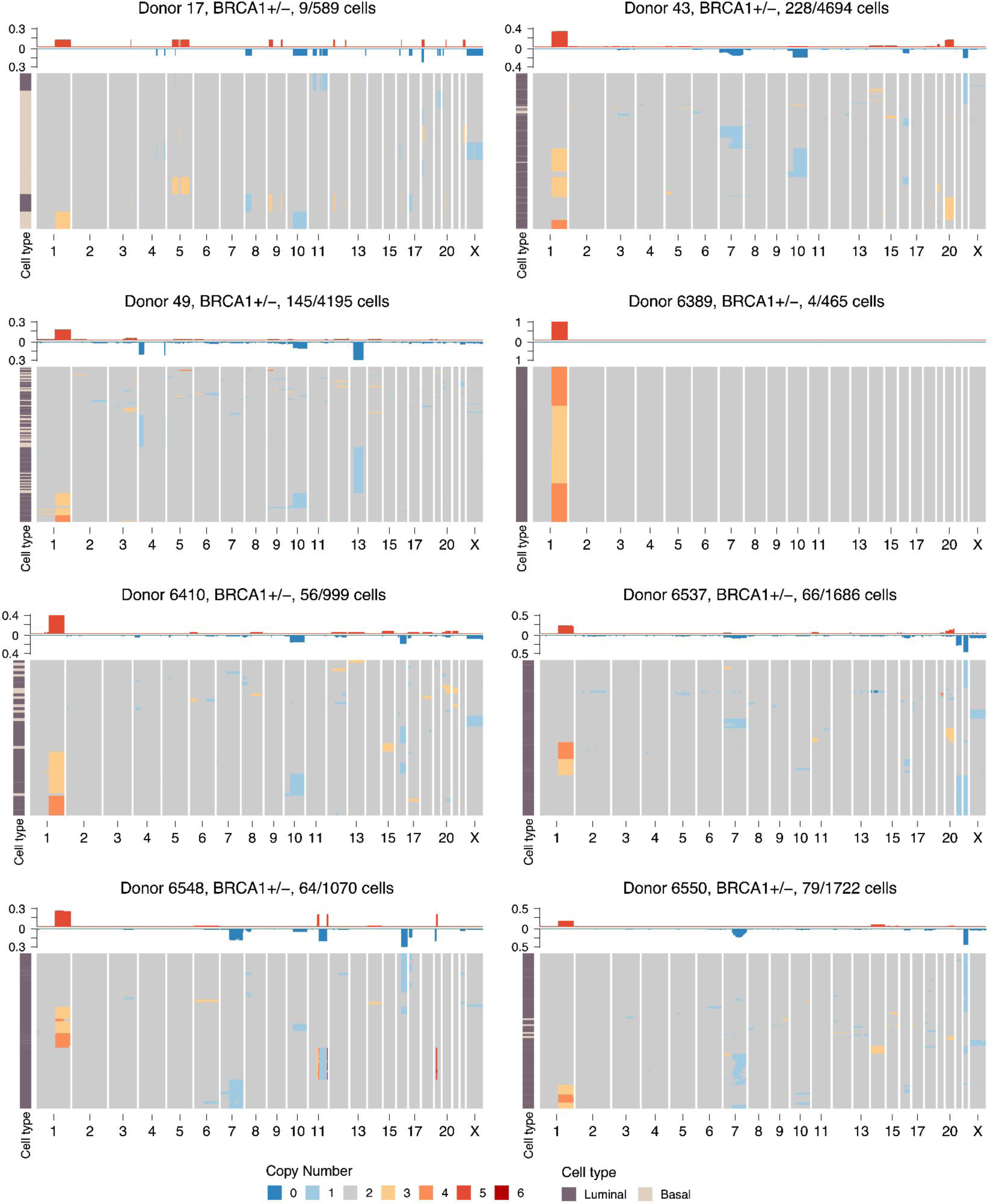
Heatmap of aneuploid cells from BRCA1 donors, title shows donor name, genotype and number of aneuploid cells out of total number of cells

**Supplementary Figure 1b.**
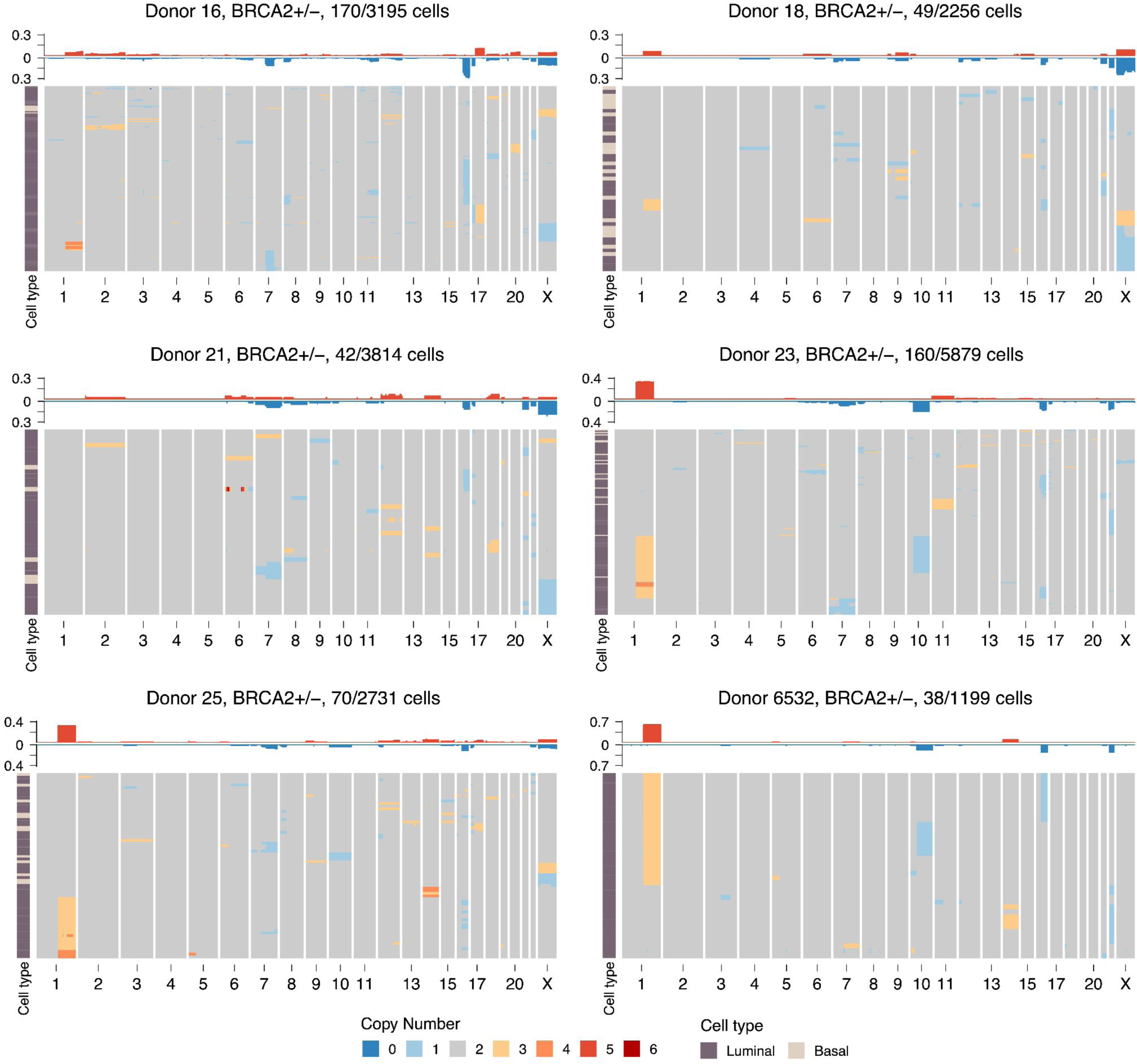
Heatmap of aneuploid cells from BRCA2 donors, title shows donor name, genotype and number of aneuploid cells out of total number of cells

**Supplementary Figure 1c.**
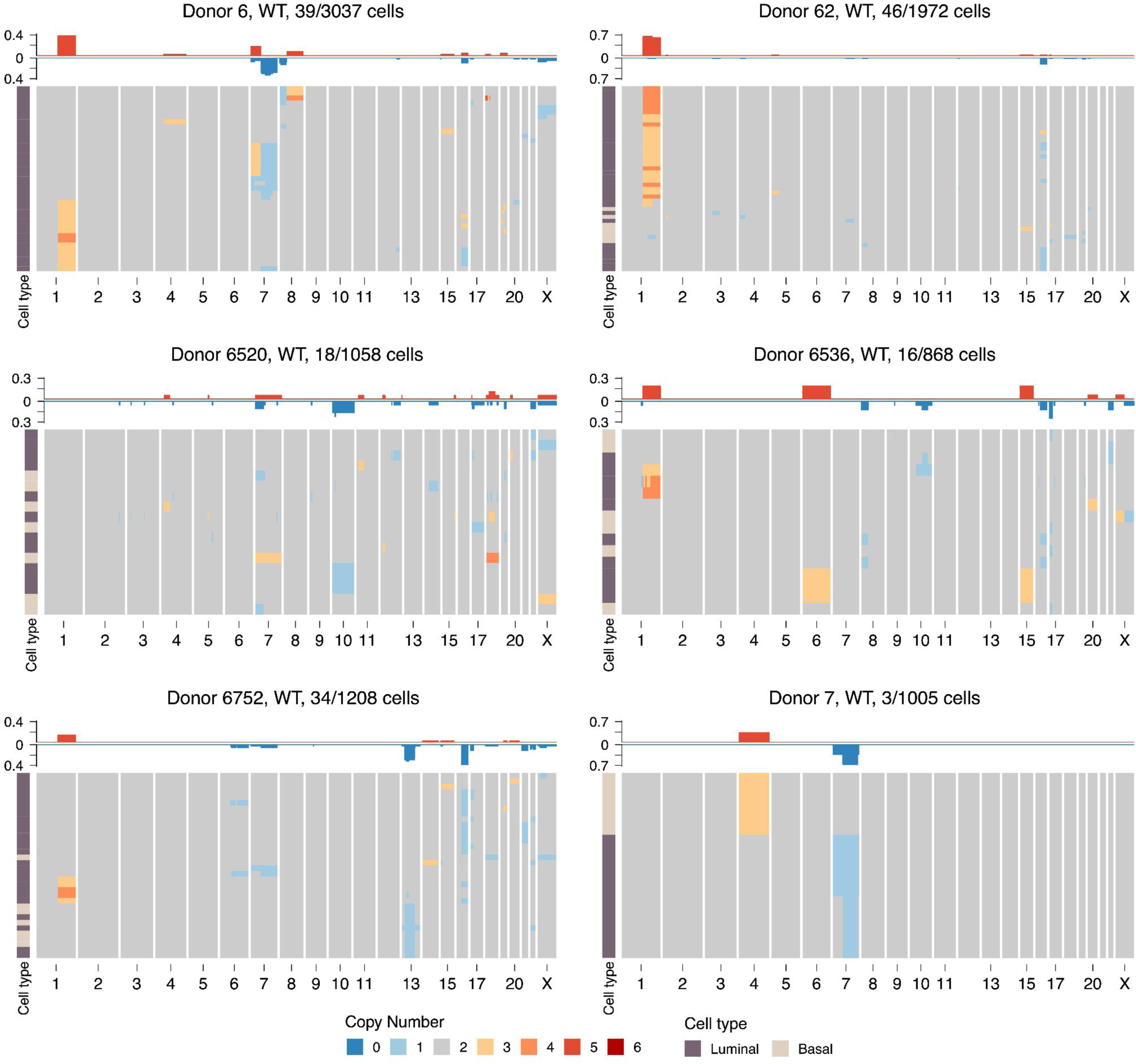
Heatmap of aneuploid cells from WT donors, title shows donor name, genotype and number of aneuploid cells out of total number of cells

**Supplementary Figure 2.**
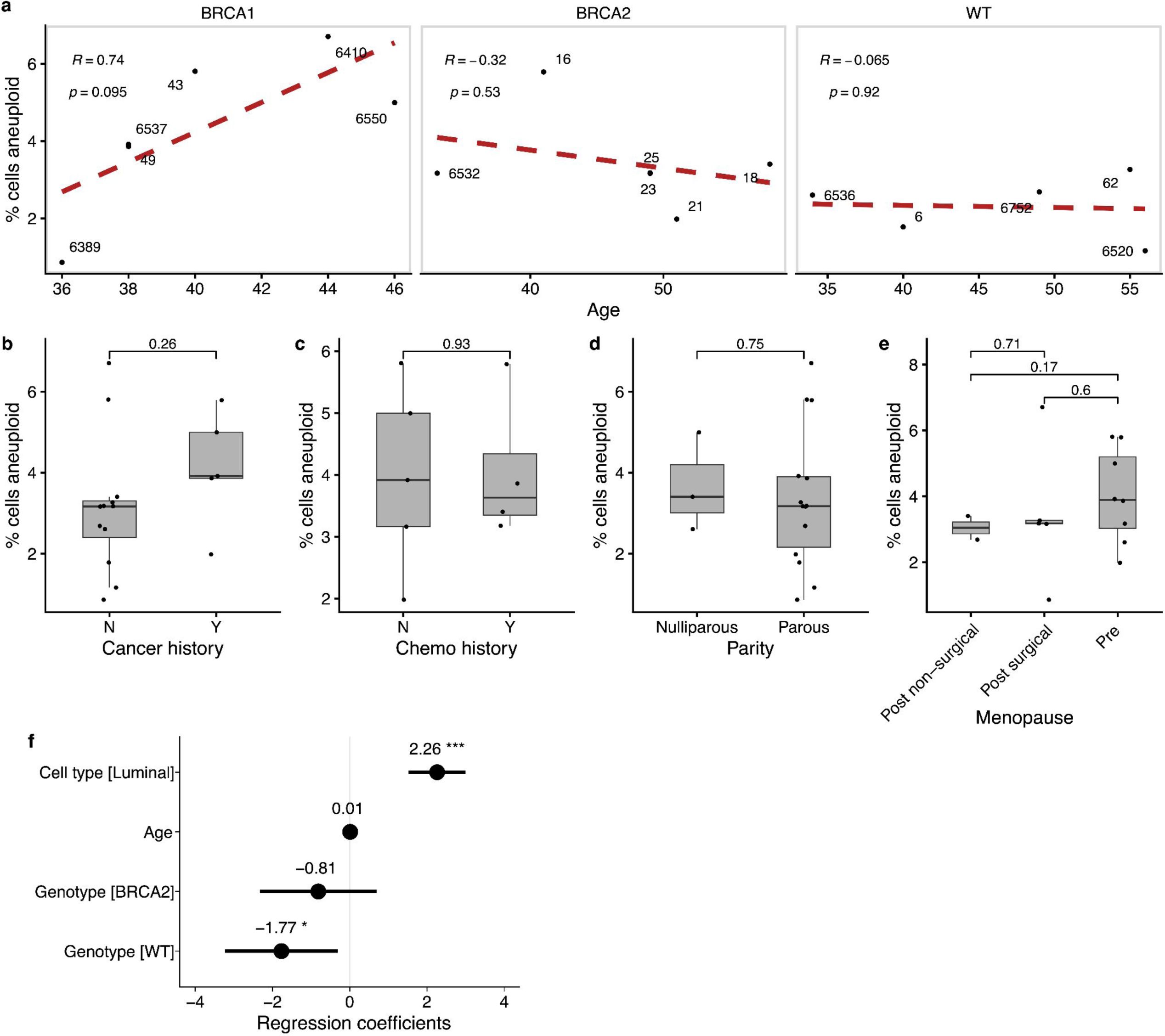
**a)** Scatter plot of % cells aneuploid vs age stratified by genotype. Red dashed lines is the linear regression line. Inset text shows correlation coefficient and p=value. Distribution of % cells aneuploid for other clinical covariates: **b)** cancer history **c)** chemo therapy history **d)** parity **e)** menopause status **f)** Coefficients of linear multivariate mixed-model, lines show 95% confidence interval

**Supplementary Figure 3.**
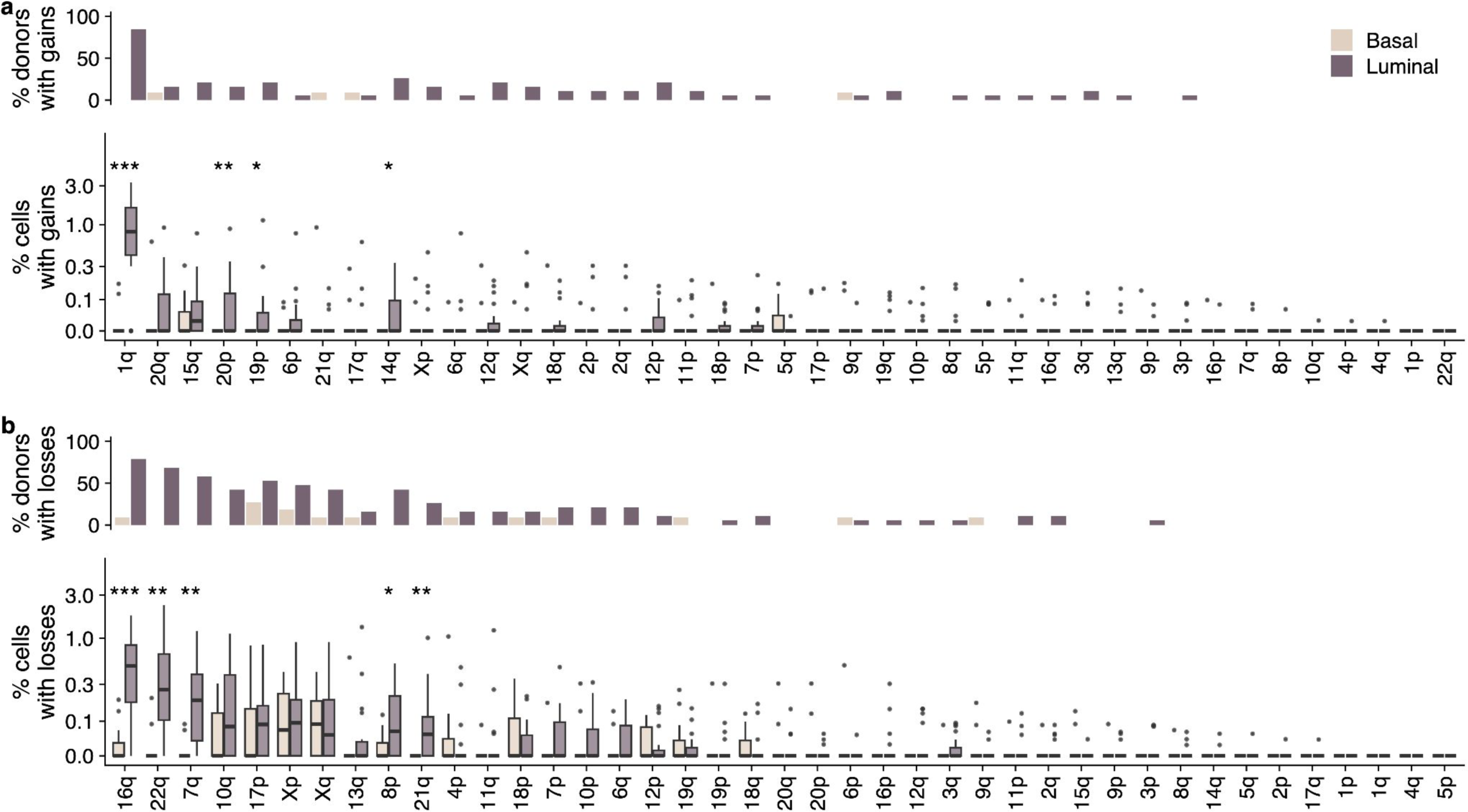
**a)** Top: % of donors that have >1 cell with chromosome arm gained per cell type. Bottom: %cells with gains per cell type, each data point is a donor. **b)** Top: % of donors that have >1 cell with chromosome arm lost per cell type. Bottom: % cells with losses per cell type, each data point is a donor.

**Supplementary Figure 4.**
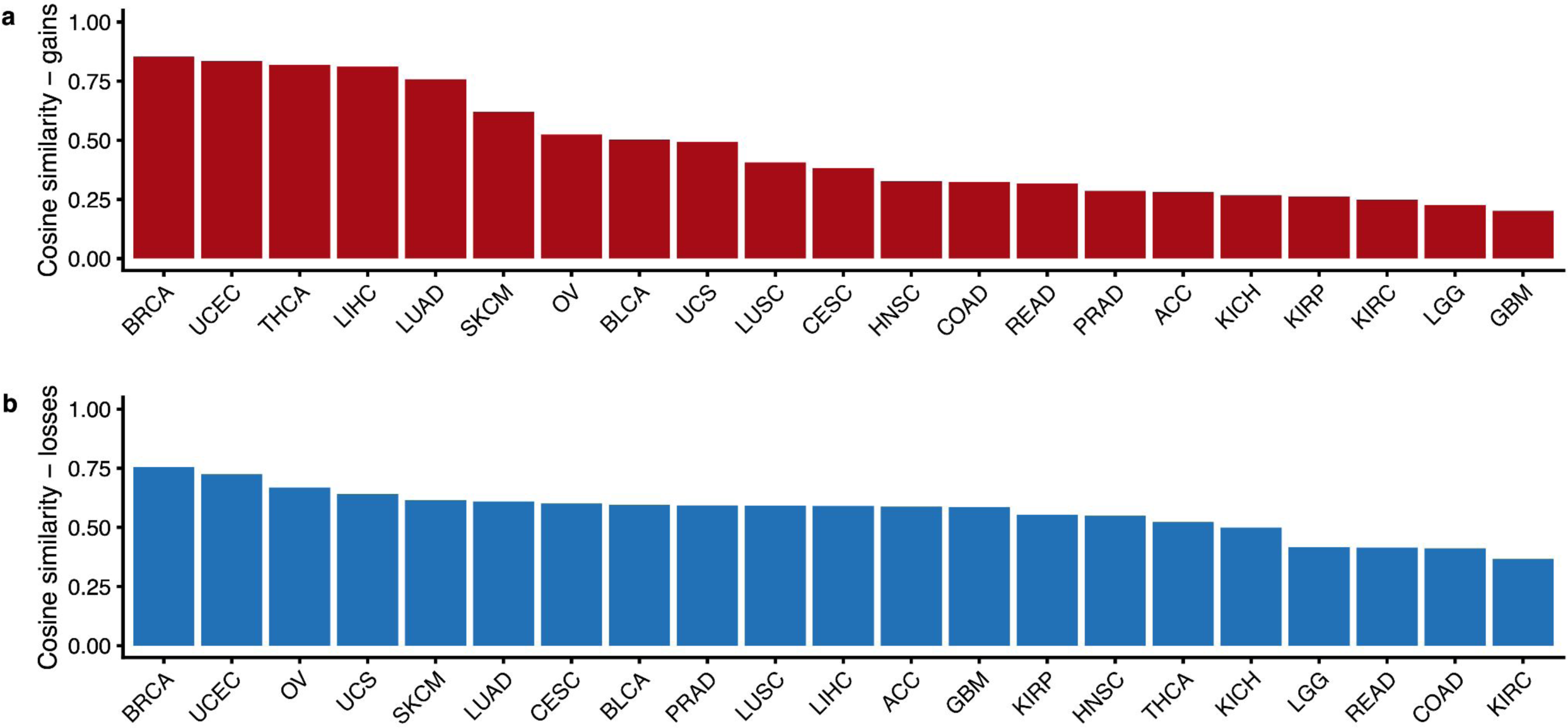
Cosine similarity between landscape of CNAs in scWGS of normal breast epithelia and TCGA subtypes for gains **a)** and losses **b)**

**Supplementary Figure 5.**
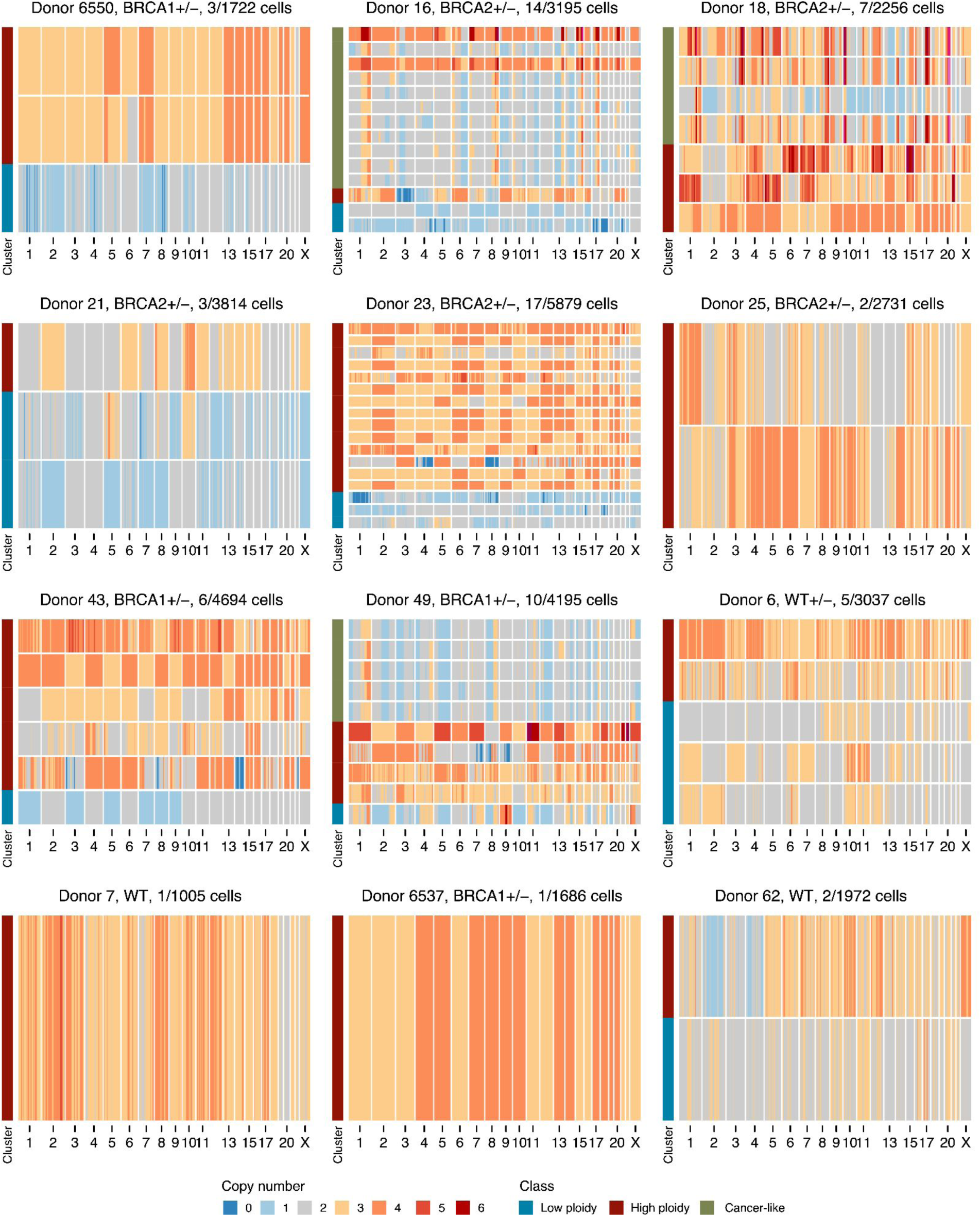
All extreme aneuploid cells per patient, title shows donor name, genotype and number of extreme aneuploid cells out of total number of cells

**Supplementary Figure 6.**
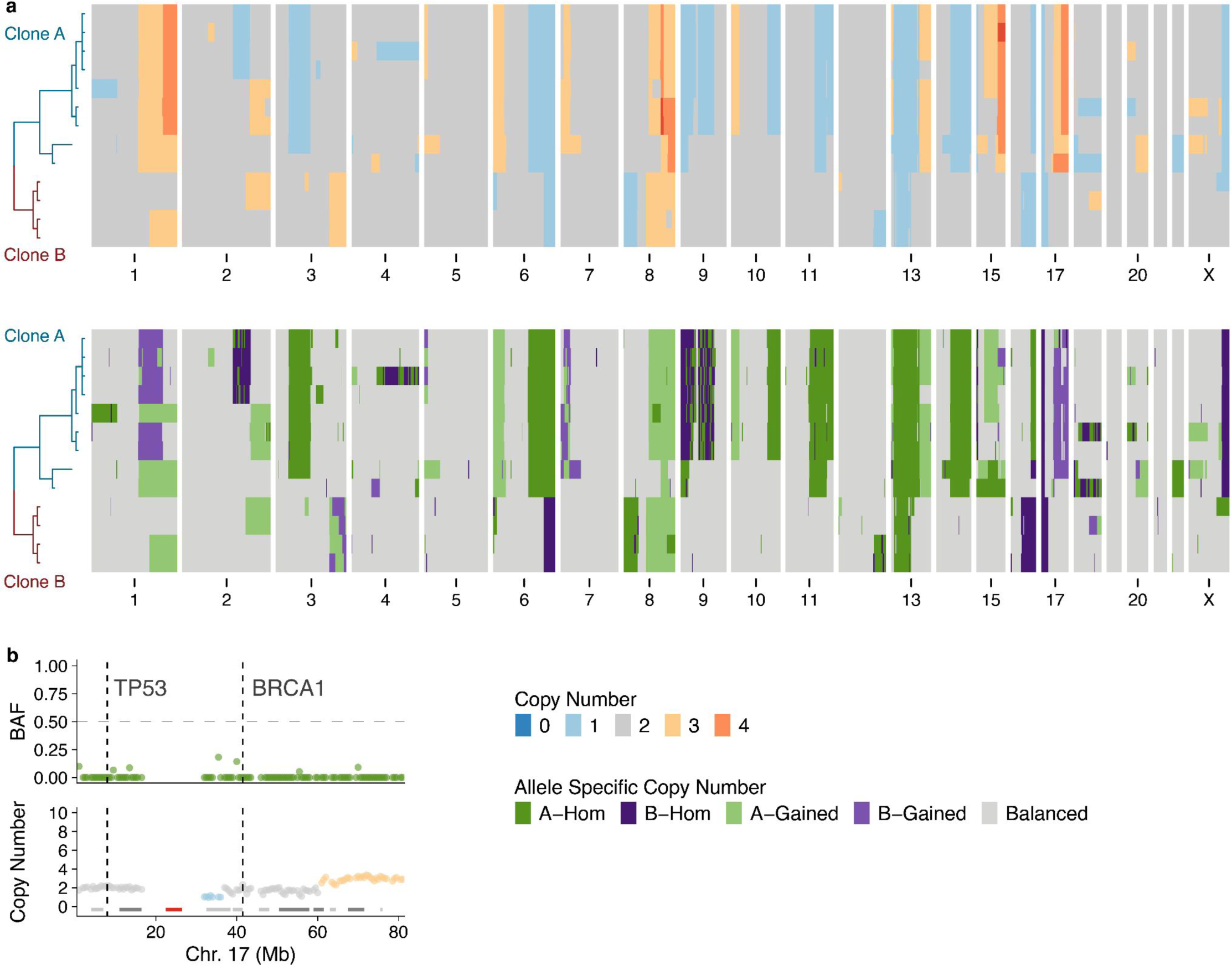
**a)** Total and allele specific copy number for the cancer-like cells in B2-16. Top shows total copy number, bottom shows allele specific copy number **b)** B-allele frequency and total copy number of chromosome 17 from donor B1-49. Location of TP53 and BRCA1 are shown with dashed lines. Data is a merged pseudobulk across the 5 cancer-like cells.

**Supplementary Figure 7.**
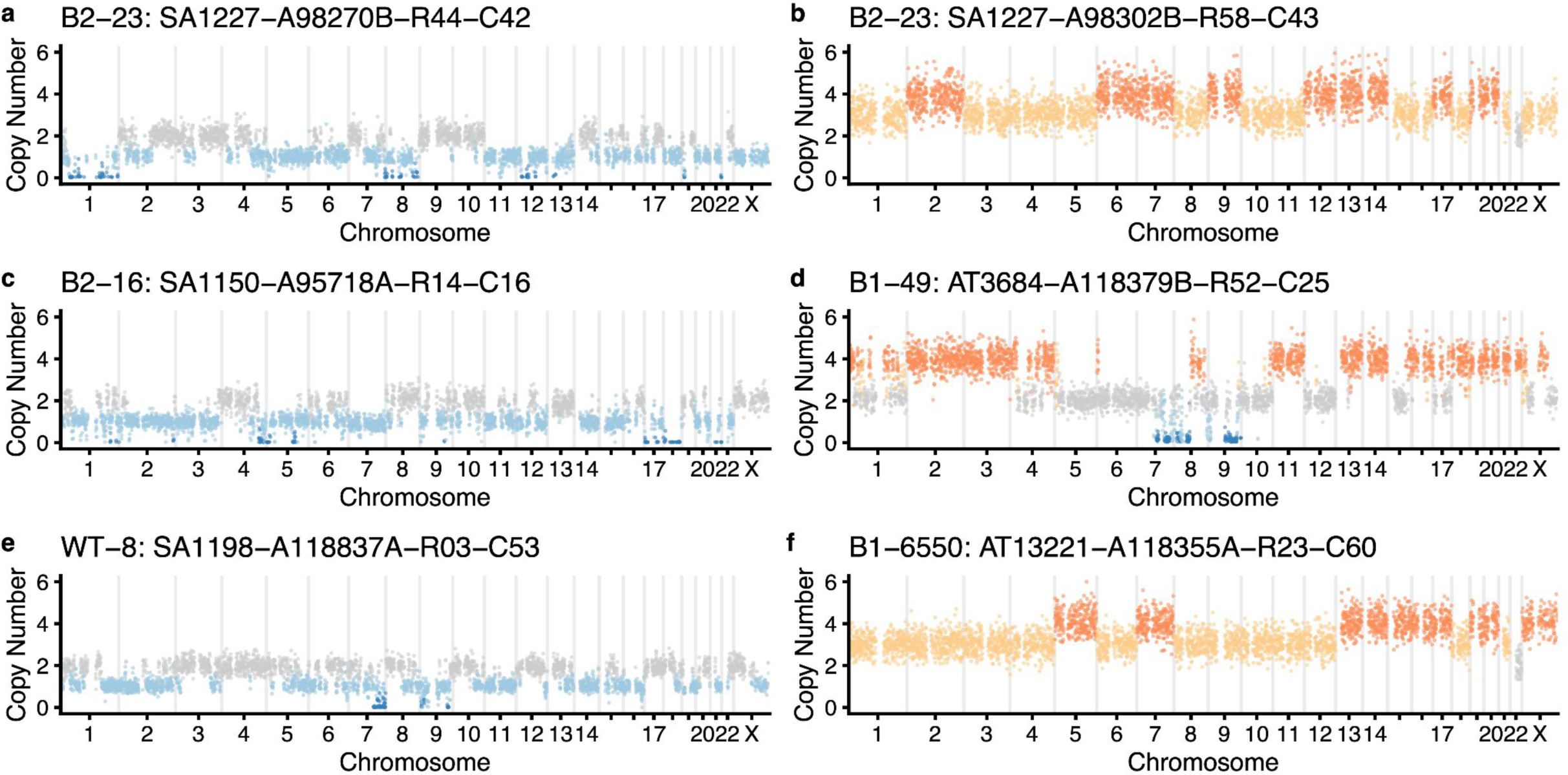
**a)-f)** Examples of extreme aneuploid genomes that are not similar to breast cancer genomes.

**Supplementary Figure 8.**
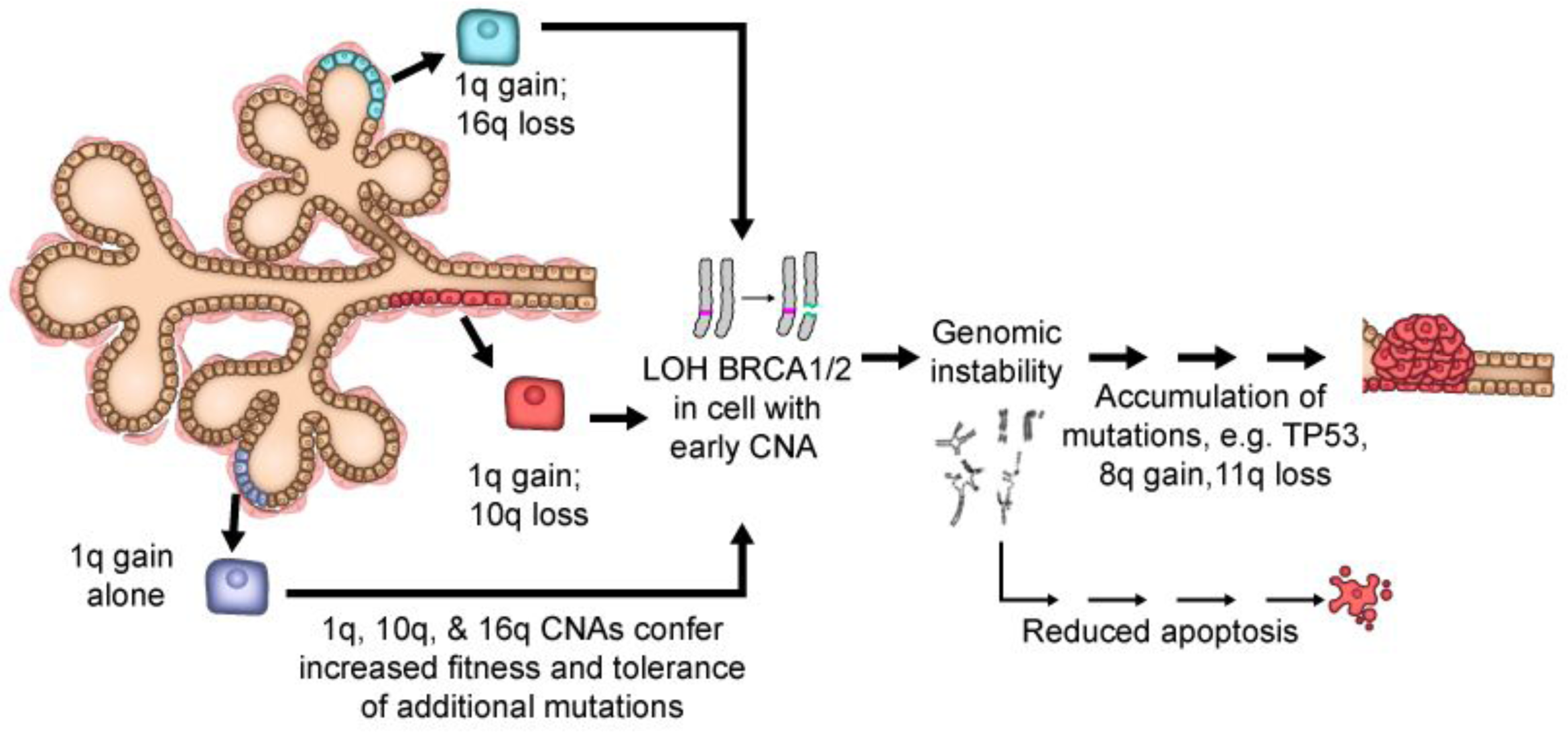
**I**n the proposed model, CNAs that accumulate in normal breast tissues (e.g. 1q gain and 10q or 16q loss) would enhance the fitness of the luminal epithelial cells. In BRCA1/2 mutation carriers, where inactivation of the wild-type (WT) copy of BRCA1/2 leads to defective DNA repair, genomic instability, and apoptosis, luminal cells carrying these CNAs would be more tolerant of these stresses, thus allowing the homologous-recombination defective mutant cells to expand, acquire oncogenic mutations, and ultimately progress to cancer.

